# Proton wires mediate the optical signal for ArcLight-type Genetically Encoded Voltage Indicators

**DOI:** 10.1101/2020.10.06.328245

**Authors:** B.E Kang, L. M. Leong, Y. Kim, K. Miyazaki, W. N. Ross, B. J. Baker

## Abstract

The genetically encoded voltage indicators, ArcLight and its derivatives, mediate voltage dependent optical signals by intermolecular, electrostatic interactions between neighboring fluorescent proteins (FPs) via proton wires. A random mutagenesis event placed a negative charge on the exterior of the FP resulting in a greater than 10-fold improvement of the voltage-dependent optical signal. Repositioning this negative charge on the exterior of the FP reversed the polarity of voltage-dependent optical signals suggesting the presence of ‘hot spots’ capable of interacting with the negative charge on a neighboring FP thereby changing the fluorescent output. To explore the potential effect on the chromophore state, voltage-clamp fluorometry was performed with alternating excitation at 390 nm followed by excitation at 470 nm resulting in several mutants exhibiting voltage-dependent, ratiometric optical signals of opposing polarities. However, the kinetics, voltage ranges, and optimal FP fusion sites were different depending on the wavelength of excitation. These results suggest that the FP has external, electrostatic pathways capable of quenching fluorescence that are wavelength specific. ArcLight-derived GEVIs may therefore offer a novel way to map how conditions external to the β-can structure can affect the fluorescence of the chromophore and transiently manipulate those pathways via conformational changes mediated by whole cell voltage clamp.

**Statement of Significance:** ArcLight-type GEVIs utilize proton pathways that send charge information outside of the FP to the internal chromophore enabling voltage induced conformational changes to affect fluorescence. These pathways are excitation wavelength specific suggesting that different external positions affect the protonated and deprotonated states of the chromophore.

## Introduction

Proton wires enable the long range transfer of protons through a protein via a chain of hydrogen bonds (1,2). This chain of hydrogen bonds restricts the proton transfer along closely spaced oxygen, nitrogen, or sulfur atoms but also allows for the rapid transfer of charge. The addition of a proton at one end of the proton wire results in the discharge of a different proton at the other end which accounts for the transfer of protons through the gramicidin A channel at a rate nearly 8 times faster than water molecules (3). This rapid transfer of charge can also affect the fluorescence of a protein since the protonation state of the chromophore affects the wavelength that excites the protein (4-6). Here, we report evidence that the ArcLight family of genetically encoded voltage indicators (GEVIs) modulate their fluorescence via intermolecular electrostatic interactions implicating the modulation of proton wires in the fluorescence protein (FP).

GEVIs convert changes in membrane potential into changes in fluorescence intensity allowing simultaneous measurements at multiple locations from subcellular regions like the endoplasmic reticulum of a neuron (7) all the way to population signals in neuronal circuits (8,9). GEVIs can be mutated to change the speed of the voltage response (10,11), the size of the optical signal (12-14), the voltage range of the optical signal (15), or the photo-physical properties of the probe (16-18). Further improvement of GEVIs would benefit from a better understanding of the mechanism(s) mediating the voltage-induced fluorescence change.

The GEVI, ArcLight, was the result of a spontaneous mutation to the FP domain (13). This spontaneous mutation introduced a negative charge to the exterior of the β-can structure of the FP resulting in a 15-fold improvement in the voltage-dependent optical signal. ArcLight consists of an FP in the cytoplasm fused to a classical four trans-membrane, voltage-sensing domain (VSD) (19) that resides in the plasma membrane enabling the GEVI to optically report changes in membrane potential (Figure 1). How alterations to the membrane potential mediate changes in fluorescence intensities, or why the addition of a negative charge outside of the voltage field in the cytoplasmic domain of the GEVI resulted in a dramatic improvement in the optical signal have been difficult to explain.

**Figure 1.**
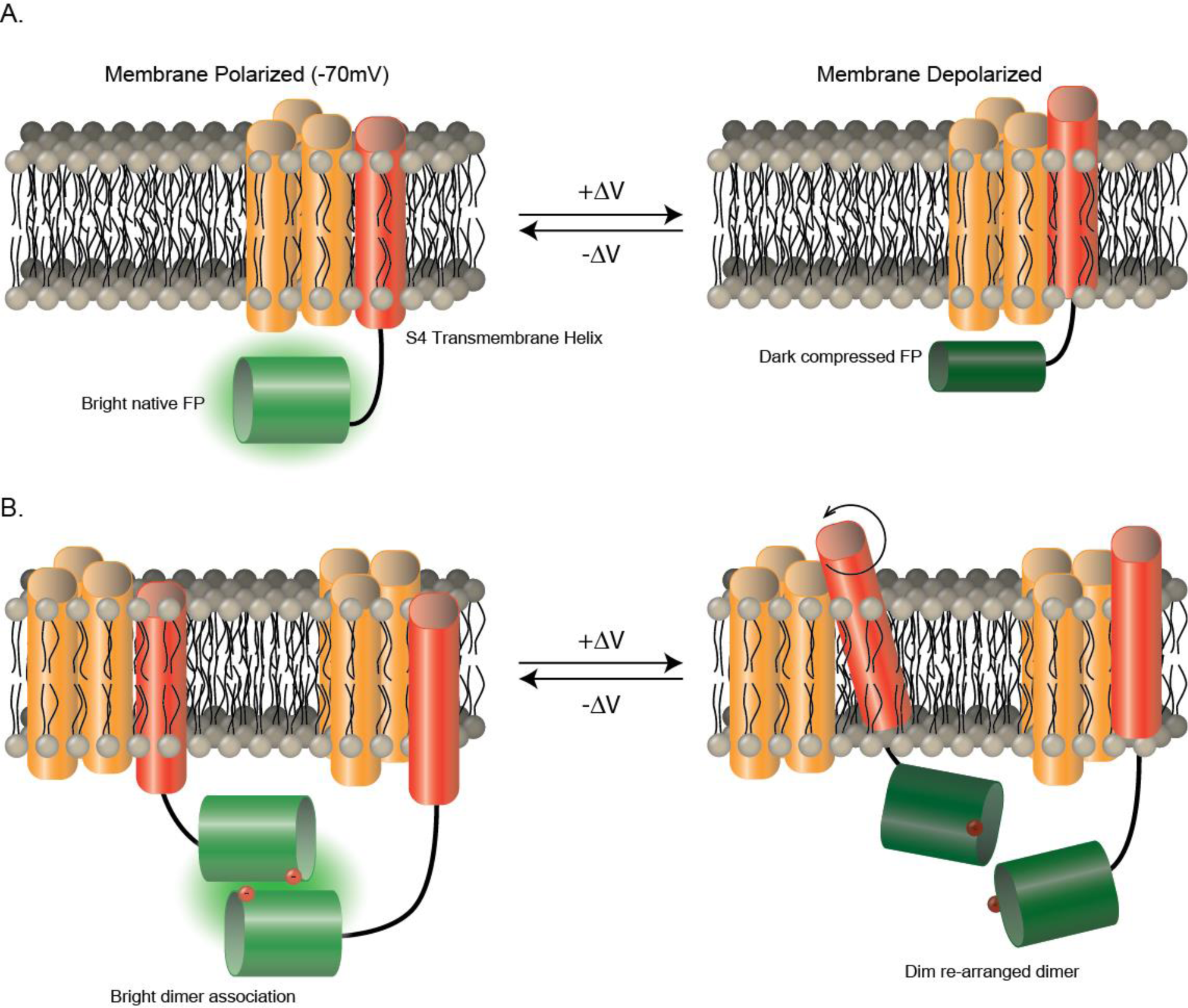
Two models for voltage-mediated fluorescence change of ArcLight-type GEVIs. A. The compression model modified from Simine et al., 2018(20) suggesting a depolarization of the plasma membrane potential (+ΔV) distorts the β-can structure causing a decrease in fluorescence. B. The dimerization model suggesting that the repositioning of a negative charge in relationship to a neighboring chromophore causes a change in fluorescence in response to voltage.

One potential mechanism suggested that physical compression of the FP domain against the plasma membrane caused the dimming of fluorescence for ArcLight in response to depolarization of the plasma membrane (Figure 1A) (20). While that report demonstrated that compression of the β-can structure could alter the electrostatic environment of the chromophore for ArcLight, the compression mechanism does not explain the requirement of an external negative charge on the FP at position 227 (A227D - numbering throughout refers to the amino acid positon in the FP) (13,16), the increase in fluorescence during hyperpolarization of the plasma membrane, or the need for the FP to be pH-sensitive (16).

An alternate model describing the mechanism of ArcLight’s voltage-dependent optical response involves an intermolecular interaction of ArcLight via the FP domain. Introduction of mutations that favor the monomeric form of the FP in ArcLight reduced the voltage-dependent optical signal by at least 70% (21) suggesting the hypothesis that conformational changes in the VSD move the external negative charge along the β-can of a neighboring FP altering the environment the chromophore (Figure 1B).

In this report we present evidence that suggests the negative charge on the exterior of the β-can mediates voltage-dependent optical signals via electrostatic interactions along proton wires (PW) on neighboring probes. Proton wires allow the shuttling of charge via defined hydrogen-bond networks. Repositioning the negative charge along the outside of the β-barrel structure of the FP resulted in the polarity of the optical response being reversed when the membrane potential was manipulated. The position of the external negative charge determined whether the fluorescence became brighter or dimmer in response to depolarization of the plasma membrane suggesting an electrostatic interaction between associated FP domains mediating the voltage-dependent optical signal. Since the FP for ArcLight is pH-sensitive, these results support the scenario that the position of the negative charge in relation to a neighboring chromophore simulates a pH-like effect on the fluorescence of the GEVI with the speed of the response now dependent upon the movement of the VSD responding to changes in membrane potential. This electrostatic interaction of ArcLight-type GEVIs offers the potential to improve the magnitude of the voltage-dependent fluorescent response as well as provide insights into how environmental changes to the exterior of the β-can structure can affect the internal chromophore of the FP.

## Material and Methods

### Subcloning

The Triple Mutant construct is described in Piao et al., 2015(10). Point mutations were made through directed mutagenesis through PCR. Oligos were synthesized (Cosmogenetech, Korea) and PCR was performed to make the target insert (BioRad). Inserts were cloned into pcDNA3.1 Hygro+ vector using restriction enzymes (NEB). Completed constructs were verified by sequencing (Cosmogenetech, Korea; Bionics, Korea).

### GEVI Expression

HEK 293 cells were maintained in DMEM supplemented with 10% v/v FBS (Gibco) at 37° C (5% CO_2_) and were seeded onto poly-L-lysine (Sigma) coated #0 glass coverslips (Ted Pella). Transfections were performed at a cell confluency of ∼70% with lipofectamine 2000 (Invitrogen). Cells were usually tested 14 hours post transfection.

### Patch clamp fluorometry

Patch clamp experiments were performed on an inverted microscope (Olympus). The chamber that held the seeded glass coverslips had a constant flow of bath solution heated to 35° C. The bath solution contained 150 mM NaCl, 4 mM KCl, 2 mM CaCl_2_,1 mM MgCl_2_, and 5 mM D-glucose and was buffered with 5 mM HEPES at pH 7.4 with NaOH. A 75 watt xenon arc lamp (Cairn) provided the light for the excitation of fluorophores. Two filter cubes were used, one comprised of an excitation filter at 472nm (FF02-472/30-25), a dichroic mirror at 495nm (FF495-Di03-25×36), and a long pass emission filter at 496 (FF01-496/LP-25). The second filter cube used the same setup except for the excitation filter which was optimized for 390nm light (FF01-386/26-25) (Semrock). The pipette electrodes were pulled from capillary tubing (World Precision Instruments) on pipette puller (Sutter Instruments) to a resistance of 3-5 MOhm. The pipette solution contained 120 mM K-aspartate, 4 mM NaCl, 4 mM MgCl_2_, 1 mM CaCl_2_, 10 mM EGTA, 3 mM Na_2_ATP and buffered with 5 mM HEPES at pH 7.2. The HEK cells were clamped and their membrane potential manipulated using a HEKA EPC10 amplifier (HEKA). An Optem Zoom System demagnifier (Qioptic) connected the fast imaging camera to the c-mount output of the microscope. The demagnified image was projected onto the 80 x 80 CCD chip of the NeuroCCD camera which was used to image fluorescence at frame rate of 1KHz (RedShirtImaging). The entire system was mounted on an air table to reduce the vibrational noise.

### Data Analysis

Data acquisition was controlled through the Neuroplex program (RedShirtImaging) which was also the platform from which we exported the ascii data for further analysis using the Origin software package (Origin Labs). Traces were averages of 4 trials and any off-line filtering is indicated in the figures.

## Results

### The position of the negative charge on the outside of the fluorescent protein determines the polarity of the voltage-dependent optical signal

The spontaneous A227D mutation to the FP of ArcLight that was responsible for a 15-fold improvement in the voltage-dependent optical signal resides on the 11^th^ β-strand with the side chain external to the b–can structure (13). The role of this external negative charge was examined by performing an aspartic acid mutagenesis scan along the external residues of the 11^th^ β-strand (Figure 2A) of an ArcLight-derived GEVI, Triple Mutant (TM), which has a longer linker segment of amino acids between the VSD and the FP and exhibits faster kinetics (10). Both constructs use the pH-sensitive FP, Super Ecliptic pHlorin (SEpH) (22,23), to optically report changes in membrane potential.

**Figure 2.**
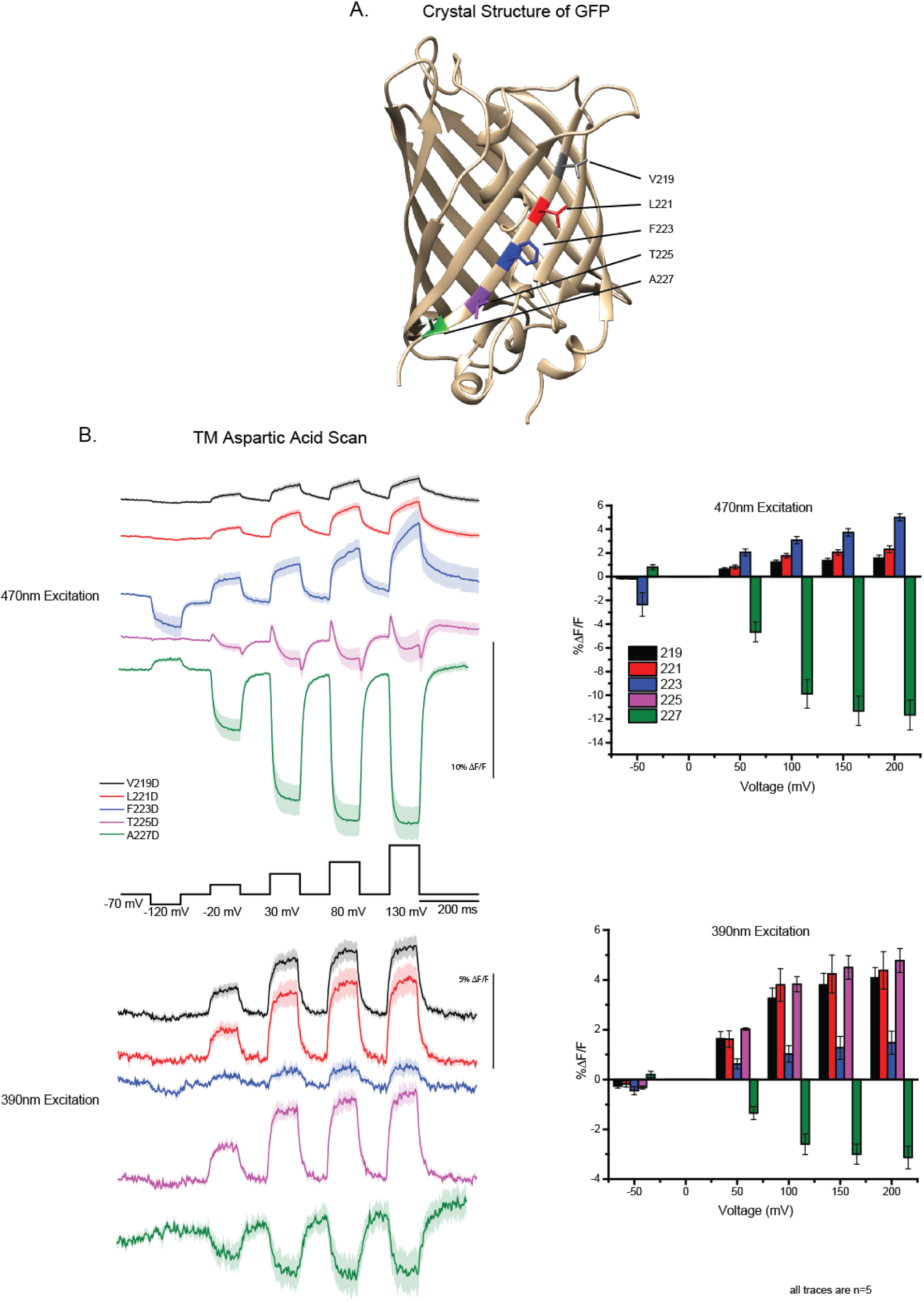
The position of the external, negative charge along the 11^th^ β-strand of the fluorescent protein determines the polarity of the voltage-dependent optical signal. A. The crystal structure of GFP from Ormö et al, (1996) (24) showing the position of the external residues along the 11^th^ β-strand. The structure was annotated using the Chimera program (25) B. Optical traces from HEK cells expressing a GEVI with the negative charge at the A227 position (A227D - green), the T225 position (T225D - purple), the F223 position (F223D - blue), the L221 position (L221D – red), or the V219 position (V219D – black). Upper traces are from experiments using 470 nm excitation light. Lower traces are from experiments using 390 nm excitation light. The command voltage of the whole-cell voltage clamp protocol is shown in black (holding potential was -70 mV). All traces were filtered offline with a Butterfield low pass 100 Hz filter.

Mutant constructs consisting of a single external negative charge at different positions along the 11^th^ β-strand were expressed in HEK 293 cells while the plasma membrane potential was manipulated via whole-cell voltage clamp (Figure 2A). When the aspartic acid residue was at the original 227 position in the FP (A227D), the optical signal got brighter upon hyperpolarization of the plasma membrane and dimmer during depolarization (Figure 2B). When the negative charge was moved to the 225 position (A227/T225D – the next residue on the β-strand with its side chain external to the β-can structure), the voltage response was quite different. The hyperpolarizing signal was nearly eliminated while the depolarizing signal consisted of at least two components with opposing polarities; there was a rapid, initial increase in fluorescence followed by a slower decrease in fluorescence. Repositioning the negative charge to the next external residue (A227/T225/F223D) resulted in the complete reversal of the polarity for the optical signal compared to the A227D construct. The hyperpolarization response got dimmer while the depolarization response got brighter. Moving the negative charge further up the β-strand to positions L221D or V219D showed similar patterns to the F223D construct albeit reduced in signal size. These results demonstrate that the position of the negative charge determines the polarity of the fluorescent response of the GEVI in response to membrane potential changes. The position of the external negative charge also affected the polarity of the voltage-dependent optical signal when TM was excited at 390 nm (Figure 2B). TM also uses the FP, SEpH, which is pH-sensitive and contains the S65T mutation to reduce the 390 nm excitation peak (22,23). Despite the S65T mutation in the FP, the aspartic acid scan constructs were capable of yielding a voltage-dependent optical signal when excited at 390 nm that also reversed orientation depending on the position of the external negative charge (Figure 2B).

### Introduction of the T65S mutation to the FP domain inverts the polarity of the optical response for TM but not ArcLight

The observation that the position of the external negative charge in the FP domain determined the polarity of the voltage-dependent optical signal suggested the possibility that the negative charge was affecting the protonation state of the chromophore. Green Fluorescent Protein (GFP) has two excitation peaks, one at 390 nm for the protonated, neutral form of the chromophore (A state), and the other at 470 nm for the anionic form of the chromophore (B state) (4-6). The ability of TM to exhibit a voltage-dependent optical signal that changed polarities when excited at 390 nm was surprising since ArcLight was previously shown to give a voltage-dependent optical signal that only decreased in fluorescence intensity when excited with 400 nm light (17).

To better understand the different behavior of these GEVIs when excited at 390 nm, we compared the ArcLight and TM constructs having the T65 version of the FP, SEpH (26), to the reverse-engineered constructs with improved 390 nm excitation (the Ecliptic pHlorin version of the FP) (22), designated epArcLight T65S and epTM T65S (Figure 3). HEK 293 cells expressing these constructs were imaged under voltage clamp fluorometry with alternate trials consisting of excitation at 390 nm followed by excitation at 470 nm on the same cell.

**Figure 3.**
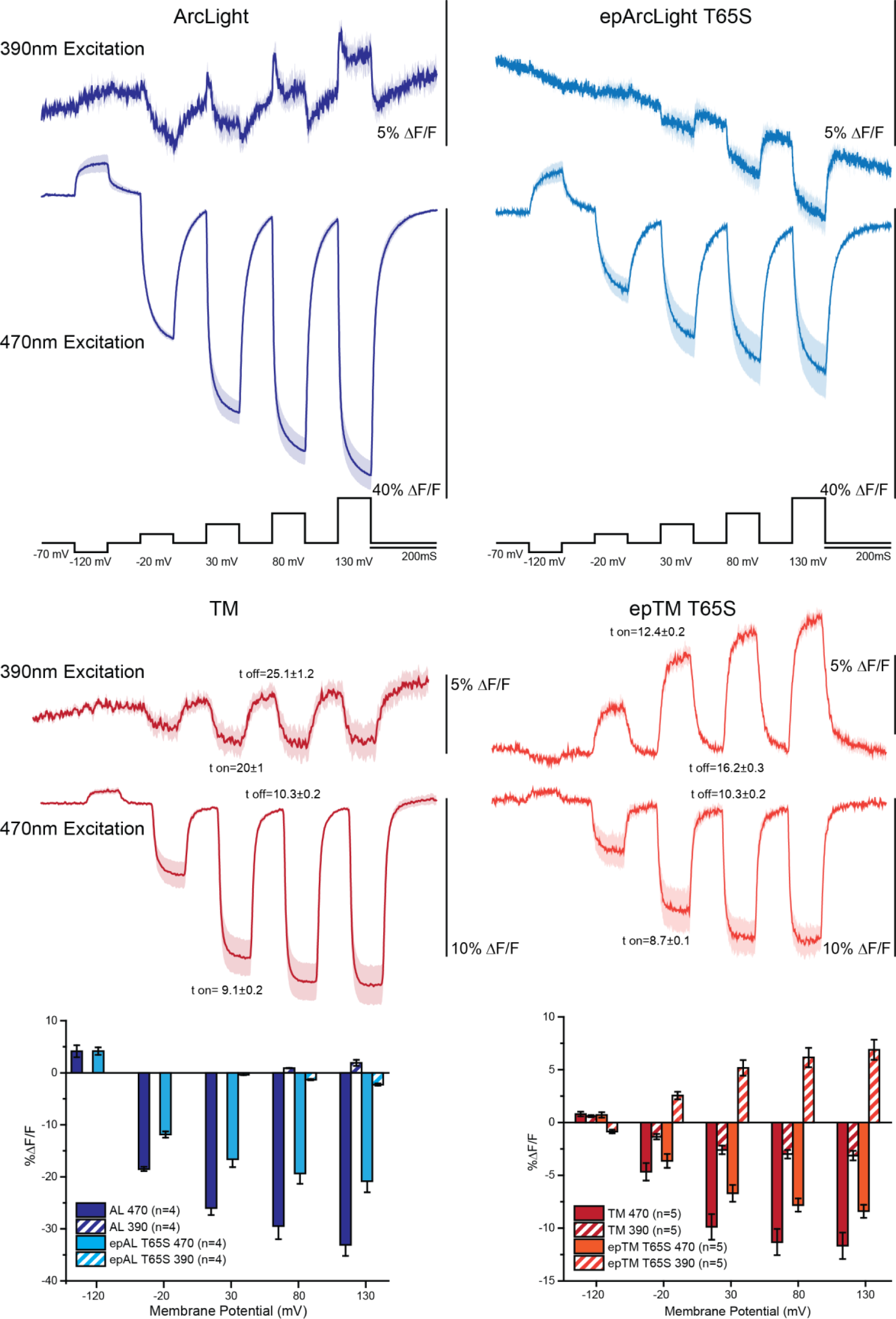
Excitation state dependence of anti-correlated fluorescence change for the GEVI, epTM T65S. Traces depict voltage-dependent optical signals from HEK cells expressing ArcLight – T65 (dark blue), epArcLight - T65S (light blue), TM - T65 (dark red), or epTM - T65S (light red). The command voltage is shown in black. For each construct, optical signalswere recorded using either 390 nm excitation light (top traces) or 470 nm excitation light (bottom traces). Shaded regions of the traces and error bars in the graphs are both standard error of the mean. All traces were filtered offline with a low pass Butterfield 100 Hz filter. Taus are given for the 100 mV depolarization step for both TM and epTM.

ArcLight (T65) gave small, but distinct voltage-dependent signals with peculiar properties when excited with 390 nm light (figure 3A). For a 50 mV depolarization of the plasma membrane, the fluorescence dimmed slightly. For a 100 mV depolarization, there was a fast component that got brighter followed by a slower component that got dimmer. Stronger depolarizations of the plasma membrane with a 150 mV or a 200 mV step exhibited a fast, transient component that got brighter and a slower, persistent component that also increased the fluorescence of the probe when compared to the fluorescence at the holding potential of -70 mV. Introduction of the T65S mutation into the FP (epArcLight) reduced the complexity of the voltage-dependent optical signal when excited at 390 nm and did not exhibit an increase in fluorescence upon depolarization of the plasma membrane when excited at 390 nm.

TM (T65) exhibited a different response than ArcLight when excited at 390 nm (Figure 3B). Regardless of excitation wavelength, the fluorescence of TM (T65) was reduced during depolarization steps. However, introduction of the T65S mutation (epTM) inverted the polarity of the voltage-dependent optical signal when excited at 390 nm. Initially we believed the inverted polarity for the epTM T65S mutant was the result of a shift in the protonation state of the chromophore. The voltage-induced conformational change during depolarization steps favored the protonated state of the chromophore causing the fluorescence to increase when excited at 390 nm and to decrease when excited at 470 nm. However, closer inspection of the optical responses for both excitation wavelengths revealed that the kinetics and voltage ranges of the optical signals for the two wavelengths were different. The tau (t) of the response which is the time constant required to achieve 63% of the maximum response for epTM T65S at 390 nm for a 100 mV depolarization step was 12.4 ± 0.2 ms while the tau of the response at 470 nm was 8.7 ± 0.1 ms. Upon returning to the holding potential, the speed of the optical response for both wavelengths were again different, exhibiting a tau of 16.2 ± 0.3 ms when excited at 390 nm compared to a tau of 10.3 ± 0.2 ms when excited with 470 nm light (Figure 3).

The voltage range was also different depending on the wavelength of excitation. The signal size for the 390 nm recording continued to increase even for the 200 mV depolarization step while the 470 nm recording plateaued at the 150 mV step suggesting different mechanisms for different wavelengths. Indeed, the mechanism of voltage-dependent fluorescence change may be chromophore state specific as the voltage-dependent optical signals for both GEVIs containing the Ecliptic pHlorin version of the FP were reduced when excited at 470 nm (Figure 3) compared to the original versions of the GEVIs.

Even though the mechanism of the voltage-induced optical signal did not involve a simple transition in the protonation state of the chromophore, the ability of the GEVIs to yield signals when excited at different wavelengths offered a potential for the development of ratio-metric probes capable of quantitating membrane potential. We therefore attempted to optimize the voltage-dependent signal for both the 390 nm and 470 nm wavelengths using the T65S version of the FP.

### The length of the linker between the VSD and the FP affects the polarity of the optical signal when excited at 390 nm but not 470 nm

A general method for improving the optical signal of GEVIs is to vary the number of amino acids between the VSD and the FP (10,27-30). We employed this strategy in the hope of generating a ratiometric probe with a dynamic response sufficient for *in vivo* recordings. GEVIs were constructed containing the Ecliptic version of the FP since epTM T65S exhibited an inverse correlation dependent upon excitation wavelength during depolarization of the plasma membrane (Figure 3). The linker segments between the VSD and the FP domain ranged from 4 to 24 amino acids (Figure 4). HEK cells expressing these constructs of varying linker lengths were voltage clamped and imaged with alternating trials of 390 nm and 470 nm excitation light. Surprisingly, the optimal linker length differed depending on the excitation wavelength. The optimal linker length for the 470 nm excitation signal was around 7-8 amino acids while a longer linker of 15 amino acids yielded the best voltage-dependent signal for 390 nm excitation. Interestingly, the polarity of the voltage-dependent optical signal during 390 nm excitation inverted as the linker length was shortened. The fluorescence decreased upon depolarization of the plasma membrane when the linker length was less than six amino acids yet the fluorescence increased for linker lengths greater than eight. This explains why little to no signal was seen previously for ArcLight upon 390 nm excitation. The linker length for ArcLight is around 8-9 amino acids depending on the version being used which are poor locations for the FP during 390 nm excitation recordings.

**Figure 4.**
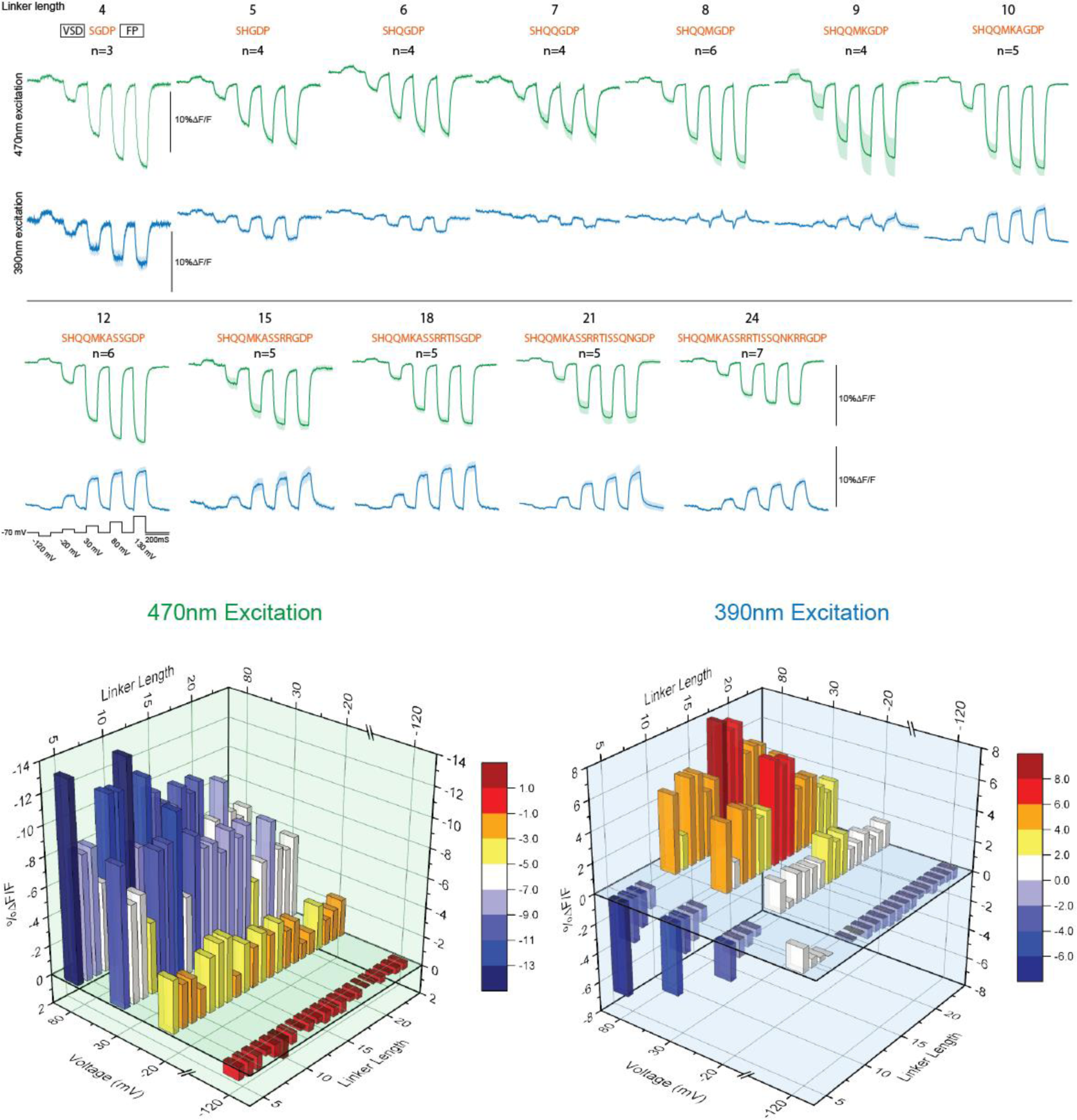
Linker length affects the voltage-dependent optical signal differently when excited at 390 nm (blue traces) than at 470 nm (green traces). Optical traces from HEK cells expressing the GEVI, epTM T65S, with varying linker lengths from four amino acids to 24 amino acids between the voltage-sensing domain (box labelled VSD) and the fluorescent protein (box labelled FP). The linker protein sequence in red is based on the *Ciona* phosphatase gene. The GDP sequence is from the Bam HI cloning site which is also included as part of the linker sequence. Numbers above indicate total number of amino acids between the VSD and the FP. The 8 and 9 linker length constructs excited at 390 nm were omitted from the bar graphs below due to the complex nature of those signals. The command voltage steps are shown in black under the 12 linker trace. All traces were filtered offline with a low pass Butterworth 100 Hz filter.

### Mutations to the E222 position can rescue the 390 nm voltage-dependent optical signal

The different optimal linker lengths for the voltage-dependent signal when excited at 390 nm compared to 470 nm suggested that the negative charge was affecting the protonated A state of the chromophore differently than the anionic B state. Different linker lengths could alter the positioning of the negative charge on the exterior of the FP in relationship to the neighboring chromophore upon movement of the VSD domain in response to voltages changes at the plasma membrane. The optimal position of the negative charge when excited at 390 nm was different than when excited at 470 nm. The ‘antennae’ or hotspot responding to the voltage-induced movement of the negative charge may be wavelength dependent.

The fluorescence of SEpH is highly sensitive to pH(22,26) indicating that a pathway or pathways exist for the proton concentration outside of the β-can structure to affect the fluorescence of the internal chromophore. Proteins have been shown to contain proton wires that allow for the rapid transfer of protons to different positons in the protein (3). Indeed, FPs are an ideal system for studying protein wires as the protonation state of the chromophore affects the fluorescent output. For instance, excitation of wildtype GFP with 390 nm light results in an excited-state proton transfer involving the protonated form of the chromophore (6,31). The E222 position has been postulated as the terminal proton acceptor in this transfer process (32,33). E222 is also near the position of the external negative charge that reverses the polarity of the voltage-dependent optical signal (Figure 2). We therefore tested four E222 mutations that varied in charge or polarity (E222Q, E222D, E222K, and E222H) in the presence of the S65 FP (390 nm excitation peak) or the T65 FP (470 nm excitation peak).

The E222Q mutants gave modest voltage-dependent optical signals when excited at 470 nm light (Figure 5). However, when excited with 390 nm light, neither the E222Q/65S nor the E222Q/65T constructs gave a voltage-dependent signal. This is consistent with the previous finding that the E222Q mutation destabilizes the protonated form of the chromophore (33).

**Figure 5.**
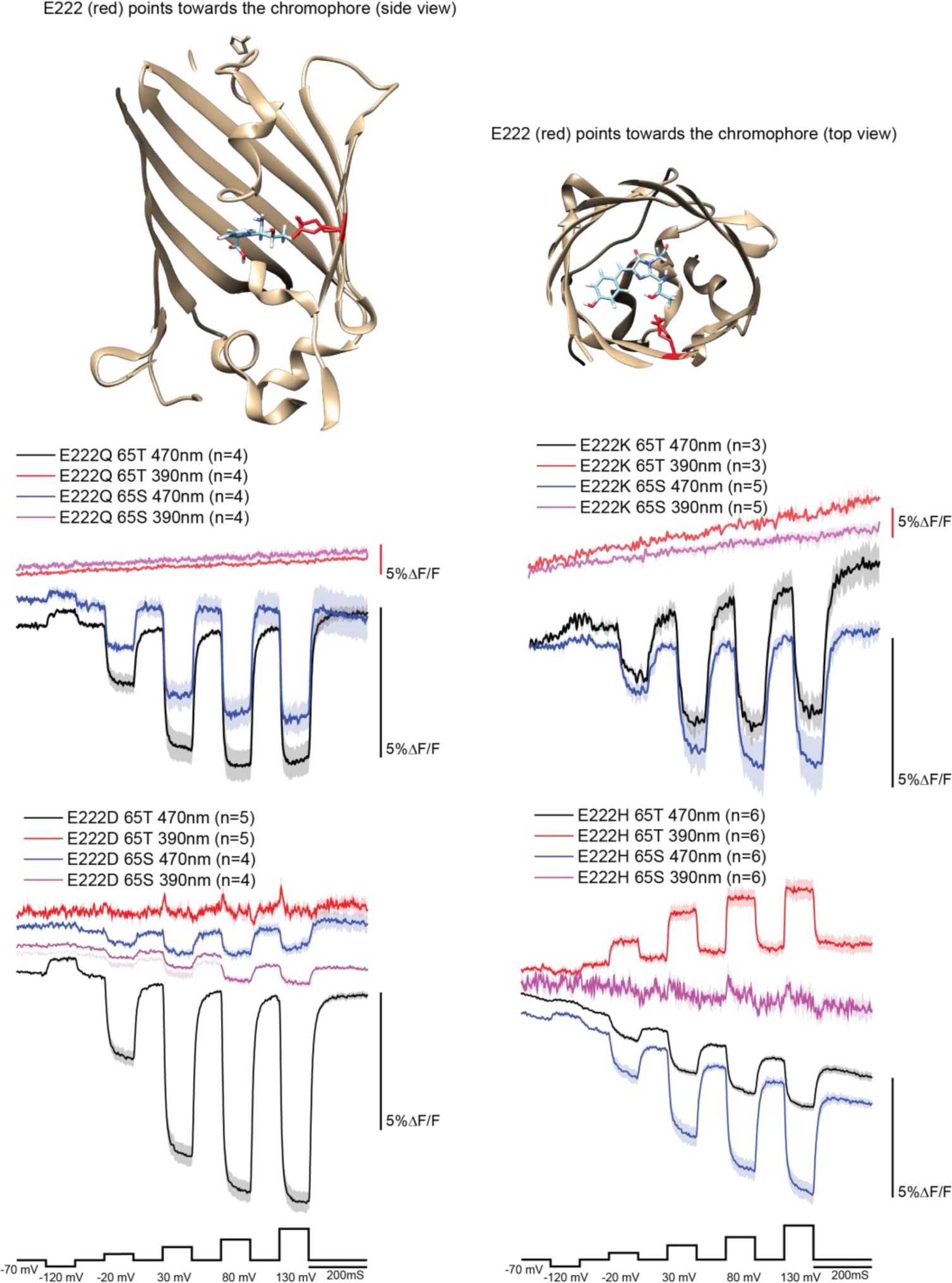
Top is the crystal structure of GFP relating the position of E222 (red) to the chromophore (blue). Below are the voltage-dependent optical traces of E222 mutants with the SEpH (65T) or the Ecliptic (65S) version of the FP expressed in HEK cells. The voltage command pulse protocol is shown at the bottom of both columns in black. Shaded regions in the traces are standard errors of the mean. All traces were filtered offline with a low pass Butterworth 100 filter.

The E222D/65T mutant response was very similar to the original GEVI, TM. The voltage-dependent optical signal was larger when excited at 470 nm than when excited at 390 nm (Figure 4). This was also the case for E222/65T construct (Figure 2). The E222D mutation in the presence of the Ecliptic S65 version of the FP exhibited a nearly 8-fold reduction in the voltage-dependent signal when excited at 470 nm (Figure 5). Interpretation of the signal for E222D/65S with 390 nm excitation with a small yet short-lived increase in fluorescence in response to depolarization of the plasma membrane is difficult.

Reversing the charge with the E222K mutations did not destroy the voltage-dependent signal when excited at 470 nm light regardless of whether there was a threonine or serine residing at position 65 (Figure 5). Excitation at 390 nm yielded very little if any fluorescence for both E222K/65T or E222K/65S constructs. As a result, no voltage-dependent signal was observed.

Like the other E222 mutations, the E222H/T65 and E222H/S65 mutants exhibited a voltage-dependent signal when excited at 470 nm (Figure 5). However, the E222H/S65 construct is the only S65 mutant with a larger signal than the T65 versions when excited at 470 nm. Surprisingly, the E222H/65T mutant also yielded an optical signal in the opposite direction when excited at 390 nm. In contrast, the E222H/65S construct yielded a very small and transient change in fluorescence when excited at 390 nm, suggesting that the E222H mutants favor the protonated form of the chromophore over the anionic chromophore. This again demonstrates that manipulating the position of an external negative charge can affect the chromophore in different ways depending on its protonation state.

## Discussion

The mechanism of voltage-mediated fluorescence change for ArcLight-type GEVIs appears to involve an electrostatic interaction between FP domains of two adjacent probes. In contrast to the structural rearrangements that might be caused by the compression of the FP onto the plasma membrane (20), the interaction of neighboring FPs creates an environment that can be altered by voltage-induced conformational changes. This FP dimerization model accounts for the effect of the A227D mutation placing a negative charge on the exterior of the β-can structure, the inverted polarity of the voltage-dependent optical signal upon hyperpolarization of the plasma membrane, the need for the FP to be pH sensitive, and the effects of varying the linker length between the VSD and the cytosolic FP.

The challenge now is to determine with which amino acids on the neighboring FP the A227D residue is interacting with. SEpH is a pH-sensitive FP that gets dimmer upon acidification of the surrounding environment (22,23,34). Initial attempts to use SEpH as the fluorescent reporter in GEVIs resulted in probes yielding a very modest change in fluorescence (1% DF/F/100 mV) upon depolarization of the plasma membrane (13), indicating that pH-sensitivity of the FP alone is not sufficient to explain the large optical signal seen for ArcLight. Fortunately, a random mutagenesis event that introduced a negatively charged amino acid on the exterior of the FP domain (A227D) resulted in a 15-fold larger voltage-dependent optical signal.

In this report, we have shown that the position of this external charge matters. Moving the negative charge along the exterior of the β-strand resulted in the reversal of the polarity of the voltage-dependent optical signal (Figure 2), suggesting that the electrostatic environment of the FP has been altered by the movement of the VSD in response to voltage. When the negative charge is at the 227 position, depolarization results in a voltage-dependent reduction of the fluorescence. When the negative charge is at positions 219, 221, or 223, the fluorescence of the GEVI gets brighter upon depolarization of the plasma membrane. This may indicate that depolarization of the plasma membrane results in the FP experiencing a more acidic environment when the negative charge is at position 227 relative to the initial environment during the holding potential. Since the lack of a negative charge on the FP results in small, voltage-dependent optical changes, we conclude that A227D is responsible for the generation of this novel environment. In agreement, introduction of monomeric favoring mutations to the FP domain of ArcLight reduced the voltage-dependent fluorescence change by over 70% (21).

One way to affect the fluorescence of the FP is to modulate the protonation state of the chromophore. GFP has two excitation peaks. The protonated state of the chromophore is excited at 390 nm while the anionic chromophore is excited at 470 nm. If the movement of the external negative charge in response to voltage affected the protonation state of a neighboring chromophore, the result would be an anti-correlated optical signal dependent upon the wavelength of excitation. Protonation of the chromophore would reduce the 470 nm excitation signal while simultaneously increasing the 390 nm excitation signal and vice versa. However, while the polarity of the voltage-dependent optical signals were anti-correlated depending on the wavelength of excitation, the speeds and voltage ranges of those optical responses were different. Further evidence suggesting that the mechanism of fluorescence change is not a simple transition in chromophore protonation state comes from the fact that the linker length separating the FP from the VSD could reverse the polarity of the voltage-dependent signal during excitation with 390 nm light but not with 470 nm excitation light (Figure 4). For constructs consisting of a linker shorter than seven amino acids, the fluorescence decreases upon depolarization of the plasma membrane regardless of whether the excitation wavelength is 390 nm or 470 nm.

Having ruled out a direct effect on the protonation state of the chromophore, the mechanism of the voltage-dependent fluorescence change most likely results from transient changes to proton wires affecting amino acids neighboring the chromophore. One indication this may be the case can be seen in the E222 mutations. E222 is the proton acceptor for the excited state proton transfer upon excitation at 390 nm. Changing E222 to aspartic acid, lysine, or glutamine abolished or greatly diminished the voltage-dependent optical signal when excited with 390 nm light. However, the E222H mutant yielded an increase in fluorescence upon depolarization of the plasma membrane when excited at 390 nm suggesting a potential rescue of the proton transfer pathway but only when the S65T mutation is present (Figure 5). Hopefully, molecular modeling simulations will reveal if this rescue is via the same proton pathway or the generation of a new one.

SEpH is capable of transmitting an environmental change in pH from the exterior of the β-can to the chromophore in the interior suggesting the presence of proton wires. There must be a route for the external environmental conditions to affect the chromophore. Indeed, a recently developed red fluorescent protein, stagRFP, was rationally designed by neutralizing an external acidic residue on the β-barrel (35). This led to a reduction in the blinking to the dark state which improved the fluorescence of the FP.

Our current hypothesis, based on the evidence presented here, is that the movement of the VSD alters a proton wire by repositioning a negative charge near the pH-sensing region of a neighboring FP. This would also explain why different linker lengths alter the voltage-dependent signal in different ways, as the pathway the negative charge of the A227D residue transversus upon movement of the VSD would be different. Figure 4 shows that the optimal linker length for the voltage-dependent signal for 390 nm excitation recordings is indeed different from that at 470 nm. If proton pathways are responsible for mediating this voltage-dependent signal, then ArcLight-type GEVIs offer the exciting possibility of differentiating/mapping these pathways by transiently altering electrostatic interactions between FP domains. The pathways could be controlled by manipulating the voltage of the plasma membrane. Indeed, by combining molecular modeling and structural data, it may be possible to map the movement of the FP in conjunction with the movement of the VSD, further improving the development of GEVIs as well as revealing how protons are moving through the FP.

## Author contributions

Bok Eum Kang designed and performed experiments, analyzed data and assisted in the writing of the manuscript. Lee Min Leong assisted with experiments involving variable linker lengths. Yoonkyung Kim assisted with experiments positioning the negative charge along the exterior of the FP. Kenichi Miyazaki assisted with the ratiometric experiments and data analysis. William N. Ross assisted with the ratiometric experiments and data analysis. Bradley J. Baker designed experiments, assisted with data analysis, and help write the manuscript.

## Acknowledgments

We thank Lawrence Cohen for critical review of the manuscript. Research reported in this publication was supported by the National Institute of Neurological Disorders And Stroke of the National Institutes of Health under Award Number U01NS099691. The content is solely the responsibility of the authors and does not necessarily represent the official views of the National Institutes of Health. This study was also funded by the Korea Institute of Science and Technology (KIST) grants 2E26190, 2E26170, 2E29180, and 2E30070.

B.E.K. designed and performed experiments, analyzed data, and wrote the manuscript. L.M.L assisted in the generation of the linker lengths constructs. Y.K. assisted in the aspartic acid scan experiments, K.M. assisted in experimental design and manuscript writing. W.N.R. assisted in experimental design and manuscript writing. B.J.B. designed experiments, analyzed data, and wrote the manuscript.

## References

1. Cukierman, S. (2003) The transfer of protons in water wires inside proteins. Frontiers in bioscience : a journal and virtual library 8, s1118–1139

2. Nagle, J. F., and Morowitz, H. J. (1978) Molecular mechanisms for proton transport in membranes.Proceedings of the National Academy of Sciences of the United States of America 75, 298–302

3. Pomes, R., and Roux, B. (1996) Structure and dynamics of a proton wire: a theoretical study of H+ translocation along the single-file water chain in the gramicidin A channel. Biophysical journal 71, 19–39

4. Chattoraj, M., King, B. A., Bublitz, G. U., and Boxer, S. G. (1996) Ultra-fast excited state dynamics in green fluorescent protein: multiple states and proton transfer. Proceedings of the National Academy of Sciences of the United States of America 93, 8362–8367

5. Agmon, N. (2005) Elementary steps in excited-state proton transfer. The journal of physical chemistry. A 109, 13–35

6. Heim, R., Prasher, D. C., and Tsien, R. Y. (1994) Wavelength mutations and posttranslational autoxidation of green fluorescent protein. Proceedings of the National Academy of Sciences of the United States of America 91, 12501–12504

7. Sepehri Rad, M., Cohen, L. B., Braubach, O., and Baker, B. J. (2018) Monitoring voltage fluctuations of intracellular membranes. Scientific Reports 8, 6911

8. Nakajima, R., and Baker, B. J. (2018) Mapping of excitatory and inhibitory postsynaptic potentials of neuronal populations in hippocampal slices using the GEVI, ArcLight. Journal of Physics D: Applied Physics 51

9. Storace, D. A., and Cohen, L. B. (2017) Measuring the olfactory bulb input-output transformation reveals a contribution to the perception of odorant concentration invariance. Nature communications 8, 81

10. Piao, H. H., Rajakumar, D., Kang, B. E., Kim, E. H., and Baker, B. J. (2015) Combinatorial mutagenesis of the voltage-sensing domain enables the optical resolution of action potentials firing at 60 Hz by a genetically encoded fluorescent sensor of membrane potential. The Journal of neuroscience : the official journal of the Society for Neuroscience 35, 372–385

11. Kralj, J. M., Douglass, A. D., Hochbaum, D. R., Maclaurin, D., and Cohen, A. E. (2012) Optical recording of action potentials in mammalian neurons using a microbial rhodopsin. Nature methods 9, 90–95

12. Lee, S., Geiller, T., Jung, A., Nakajima, R., Song, Y.-K., and Baker, B. J. (2017) Improving a genetically encoded voltage indicator by modifying the cytoplasmic charge composition. Scientific Reports 7, 8286

13. Jin, L., Han, Z., Platisa, J., Wooltorton, J. R., Cohen, L. B., and Pieribone, V. A. (2012) Single action potentials and subthreshold electrical events imaged in neurons with a fluorescent protein voltage probe. Neuron 75, 779–785

14. Chamberland, S., Yang, H. H., Pan, M. M., Evans, S. W., Guan, S., Chavarha, M., Yang, Y., Salesse, C., Wu, H., Wu, J. C., Clandinin, T. R., Toth, K., Lin, M. Z., and St-Pierre, F. (2017) Fast two-photon imaging of subcellular voltage dynamics in neuronal tissue with genetically encoded indicators. Elife 6

15. Jung, A., Rajakumar, D., Yoon, B. J., and Baker, B. J. (2017) Modulating the Voltage-sensitivity of a Genetically Encoded Voltage Indicator. Experimental neurobiology 26, 241–251

16. Han, Z., Jin, L., Chen, F., Loturco, J. J., Cohen, L. B., Bondar, A., Lazar, J., and Pieribone, V. A. (2014) Mechanistic studies of the genetically encoded fluorescent protein voltage probe ArcLight. PloS one 9, e113873

17. Platisa, J., Vasan, G., Yang, A., and Pieribone, V. A. (2017) Directed Evolution of Key Residues in Fluorescent Protein Inverses the Polarity of Voltage Sensitivity in the Genetically Encoded Indicator ArcLight. ACS Chem Neurosci 8, 513–523

18. Yi, B., Kang, B. E., Lee, S., Braubach, S., and Baker, B. J. (2018) A dimeric fluorescent protein yields a bright, red-shifted GEVI capable of population signals in brain slice. Scientific Reports 8, 15199

19. Murata, Y., Iwasaki, H., Sasaki, M., Inaba, K., and Okamura, Y. (2005) Phosphoinositide phosphatase activity coupled to an intrinsic voltage sensor. Nature 435, 1239–1243

20. Simine, L., Lammert, H., Sun, L., Onuchic, J. N., and Rossky, P. J. (2018) Fluorescent Proteins Detect Host Structural Rearrangements via Electrostatic Mechanism. Journal of the American Chemical Society 140, 1203–1206

21. Kang, B. E., and Baker, B. J. (2016) Pado, a fluorescent protein with proton channel activity can optically monitor membrane potential, intracellular pH, and map gap junctions. Sci Rep 6, 23865

22. Miesenbock, G., De Angelis, D. A., and Rothman, J. E. (1998) Visualizing secretion and synaptic transmission with pH-sensitive green fluorescent proteins. Nature 394, 192–195

23. Ng, M., Roorda, R. D., Lima, S. Q., Zemelman, B. V., Morcillo, P., and Miesenbock, G. (2002) Transmission of olfactory information between three populations of neurons in the antennal lobe of the fly. Neuron 36, 463–474

24. Ormo, M., Cubitt, A. B., Kallio, K., Gross, L. A., Tsien, R. Y., and Remington, S. J. (1996) Crystal structure of the Aequorea victoria green fluorescent protein. Science 273, 1392–1395

25. Pettersen, E. F., Goddard, T. D., Huang, C. C., Couch, G. S., Greenblatt, D. M., Meng, E. C., and Ferrin, T. E. (2004) UCSF Chimera--a visualization system for exploratory research and analysis. Journal of computational chemistry 25, 1605–1612

26. Sankaranarayanan, S., De Angelis, D., Rothman, J. E., and Ryan, T. A. (2000) The use of pHluorins for optical measurements of presynaptic activity. Biophysical journal 79, 2199–2208

27. Dimitrov, D., He, Y., Mutoh, H., Baker, B. J., Cohen, L., Akemann, W., and Knopfel, T. (2007) Engineering and characterization of an enhanced fluorescent protein voltage sensor. PloS one 2, e440

28. Jung, A., Garcia, J. E., Kim, E., Yoon, B. J., and Baker, B. J. (2015) Linker length and fusion site composition improve the optical signal of genetically encoded fluorescent voltage sensors. Neurophotonics 2, 021012

29. Yi, B., Kang, B. E., Lee, S., Braubach, S., and Baker, B. J. (2018) A dimeric fluorescent protein yields a bright, red-shifted GEVI capable of population signals in brain slice. Sci Rep 8, 15199

30. Han, Z., Jin, L., Platisa, J., Cohen, L. B., Baker, B. J., and Pieribone, V. A. (2013) Fluorescent protein voltage probes derived from ArcLight that respond to membrane voltage changes with fast kinetics. PloS one 8, e81295

31. Brejc, K., Sixma, T. K., Kitts, P. A., Kain, S. R., Tsien, R. Y., Ormo, M., and Remington, S. J. (1997) Structural basis for dual excitation and photoisomerization of the Aequorea victoria green fluorescent protein. Proceedings of the National Academy of Sciences of the United States of America 94, 2306–2311

32. Lill, M. A., and Helms, V. (2002) Proton shuttle in green fluorescent protein studied by dynamic simulations. Proceedings of the National Academy of Sciences of the United States of America 99, 2778–2781

33. Jung, G., Wiehler, J., and Zumbusch, A. (2005) The photophysics of green fluorescent protein: influence of the key amino acids at positions 65, 203, and 222. Biophysical journal 88, 1932–1947

34. Shen, Y., Rosendale, M., Campbell, R. E., and Perrais, D. (2014) pHuji, a pH-sensitive red fluorescent protein for imaging of exo- and endocytosis. The Journal of cell biology 207, 419–432

35. Mo, G. C. H., Posner, C., Rodriguez, E. A., Sun, T., and Zhang, J. (2020) A rationally enhanced red fluorescent protein expands the utility of FRET biosensors. Nature communications 11, 1848

